# Optimal Transport improves cell-cell similarity inference in single-cell omics data

**DOI:** 10.1101/2021.03.19.436159

**Authors:** Geert-Jan Huizing, Gabriel Peyré, Laura Cantini

## Abstract

The recent advent of high-throughput single-cell molecular profiling is revolutionizing biology and medicine by unveiling the diversity of cell types and states contributing to development and disease. The identification and characterization of cellular heterogeneity is typically achieved through unsupervised clustering, which crucially relies on a similarity metric.

We here propose the use of Optimal Transport (OT) as a cell-cell similarity metric for single-cell omics data. OT defines distances to compare, in a geometrically faithful way, high-dimensional data represented as probability distributions. It is thus expected to better capture complex relationships between features and produce a performance improvement over state-of-the-art metrics. To speed up computations and cope with the high-dimensionality of single-cell data, we consider the entropic regularization of the classical OT distance. We then extensively benchmark OT against state-of-the-art metrics over thirteen independent datasets, including simulated, scRNA-seq, scATAC-seq and single-cell DNA methylation data. First, we test the ability of the metrics to detect the similarity between cells belonging to the same groups (e.g. cell types, cell lines of origin). Then, we apply unsupervised clustering and test the quality of the resulting clusters.

In our in-depth evaluation, OT is found to improve cell-cell similarity inference and cell clustering in all simulated and real scRNA-seq data, while its performances are comparable with Pearson correlation in scATAC-seq and single-cell DNA methylation data. All our analyses are reproducible through the OT-scOmics Jupyter notebook available at https://github.com/ComputationalSystemsBiology/OT-scOmics.

## Introduction

Allowing the measurement of gene expression in thousands of cells in a single experiment, single-cell RNA sequencing (scRNA-seq) has unveiled the diversity of the cells constituting human tissues [1]. The possibility to assess cellular heterogeneity at a previously inaccessible resolution has profoundly impacted our understanding of development, of the immune system functioning, and of many diseases [2–4]. While scRNA-seq is now mature, the single-cell technological development has shifted to the measurement of other omics, e.g. DNA methylation, proteome, and chromatin accessibility [5,6].

A common goal in single-cell data analysis is the identification of the cell types and cell states present in a sample [7]. This is typically achieved in a data-driven fashion through unsupervised clustering [8,9]. Cells with similar transcriptional profiles are assembled into clusters, which are then annotated based on markers [8]. As a consequence, the quality of such clustering plays a critical role in the derived biological discovery. While numerous clustering algorithms have been proposed, they all rely on a similarity metric for categorizing individual cells. Popular metrics include the Euclidean and Manhattan distances, Cosine similarity and Pearson correlation [10–13].

Optimal Transport (OT) emerged in the last decade as a promising mathematical toolkit to analyze and compare high dimensional data using different variants of the Wasserstein distance [14,15]. Recently, applications of OT to biology have been proposed [16–20]. Some works use OT to compare cells in the context of trajectory inference [16] and alignment of unpaired (i.e. independently profiled) scRNA-seq and scATAC-seq data [17,18]. Others apply OT to compare genes in scRNA-seq data and perform supervised classification, such as malignant vs. normal samples [19]. Finally, we recently developed a fixed point method which combines the two approaches and performs joint optimal transport over both genes and cells [20].

Here, we propose the use of OT as a cell-cell similarity metric for single-cell data. Classical OT, as applied in Bellazzi et al. [19], requires solving a costly linear optimization problem. We here use OT with entropic regularization [21], allowing to break the curse of dimensionality and efficiently compute the OT distance on a Graphics Processing Unit (GPU) for large numbers of cells [22]. We further extensively benchmark OT against state-of-the-art metrics. We apply the different metrics to single-cell data with known groups (e.g. cell types, cell lines of origin) and we evaluate their ability to detect the similarity between cells belonging to the same group. We then apply different unsupervised clustering algorithms to the computed distance matrices and test the quality of the resulting clusters. Of note, all the tests are performed in three conditions: (i) simulated scRNA-seq data, where the effect of the number of cells and of the size and overlap of the clusters can be tested in-depth; (ii) real scRNA-seq data, profiled from cell lines and colorectal tissue; and (iii) other omics data, including scATAC-seq and single-cell DNA methylation.

All the performed analyses are reproducible using the OT-scOmics Jupyter notebook provided on GitHub (https://github.com/ComputationalSystemsBiology/OT-scOmics). Users can also employ OT-scOmics to test the various metrics on new single-cell data and to evaluate the performances of other/new metrics.

## Methods

### scRNA-seq data simulation

scRNA-seq data with 5,000 genes and three underlying clusters have been simulated using the R Bioconductor package Splatter [23]. Splatter simulates scRNA-seq data using the Splat model, built around a gamma-Poisson distribution. Different parameters can be tuned in the Splatter simulation. Details on the parameters used to run Splatter are available in Supplementary Text.

Using simulated scRNA-seq data, we can assess in-depth the influence of different factors on the quality of the inferred cell-cell similarities. We simulated five scRNA-seq datasets (Table 1) obtained by varying three main factors:

i. The number of cells constituting the scRNA-seq dataset (batchCells parameter in Splatter). Datasets with 500, 1,000 and 10,000 cells are simulated. Of note, we thereby also test whether the different metrics can be computed for large numbers of cells.
ii. The overlap of the clusters (de.prob parameter in Splatter). Overlapping and well-separated clusters are simulated varying de.prob between 0.4 and 0.7, respectively.
iii. The equal or unbalanced size of the clusters (group.prob parameter in Splatter). We set the probabilities of the clusters either equal (⅓, ⅓, ⅓) or unbalanced (0.75, 0.25, 0.05). The unbalanced case reflects the more realistic scenario of a tissue composed of a mixture of prevalent and rare cell types or states.

**Table 1.** Summary of the datasets used for our benchmark and their clustering results. In the first part of the table (“Data description”), for each dataset, we specify the name with which we denote it in the paper, the reference to its original publication, the type of data, the number of cells, the number of features after preprocessing and the number of clusters (e.g. cell types, cell lines) present in the data. In the second part of the table (“Clustering results”), we report the number of clusters, chosen by maximizing the silhouette score, with hierarchical and spectral clustering for Pearson correlation and OT distance.

| Dataset name | Data description |  |  |  |  | Clustering results |  |  |  |
| --- | --- | --- | --- | --- | --- | --- | --- | --- | --- |
|  | Data type | Reference | Number of cells | Number of features after preprocessing | Number of clusters in ground-truth | Detected number of clusters for hierarchical clustering (OT) | Detected number of clusters for hierarchical clustering (Pearson) | Detected number of clusters for spectral clustering (OT) | Detected number of clusters for spectral clustering (Pearson) |
| 500 cells | Simulated scRNA-seq data | Splatter [23] | 500 | 5000 | 3 | 3 | 3 | 3 | 3 |
| 1000 cells | Simulated scRNA-seq data | Splatter [23] | 1000 | 5000 | 3 | 3 | 3 | 3 | 3 |
| 10,000 cells | Simulated scRNA-seq data | Splatter [23] | 10000 | 5000 | 3 | 3 | 5 | 3 | 3 |
| Unbalanced clusters | Simulated scRNA-seq data | Splatter [23] | 1000 | 5000 | 3 | 3 | 3 | 3 | 3 |
| Overlapping clusters | Simulated scRNA-seq data | Splatter [23] | 1000 | 5000 | 3 | 3 | 3 | 3 | 3 |
| Liu scRNA | scRNA-seq | Liu et al. [24] | 206 | 10000 | 3 | 3 | 3 | 3 | 3 |
| Li Tumor | scRNA-seq | Li et al. [25] | 364 | 10000 | 7 | 6 | 6 | 3 | 3 |
| Li NM | scRNA-seq | Li et al. [25] | 266 | 10000 | 7 | 5 | 23 | 6 | 13 |
| Li cell lines | scRNA-seq | Li et al. [25] | 561 | 10000 | 7 | 8 | 10 | 5 | 7 |
| Liu scATAC | scATAC-seq | Liu et al. [24] | 206 | 10000 | 3 | 3 | 3 | 3 | 3 |
| scMethylation mouse | Single-cell DNA methylation | Luo et al. [27] | 3377 | 10000 | 16 | 3 | 3 | 3 | 3 |
| scMethylation human | Single-cell DNA methylation | Luo et al. [27] | 2740 | 10000 | 21 | 3 | 7 | 3 | 10 |
| Leukemia scATAC | scATAC-seq | Corces et al. [26] | 391 | 7602 | 6 | 3 | 25 | 24 | 14 |

### Single-cell omics data acquisition and preprocessing

Several publicly available single-cell omics datasets have been employed (Table 1). Only public datasets providing ground-truth labels for all the profiled cells were considered. The labels are intended to associate each cell to a specific group (e.g. cell type, cell line of origin). Of note, labels are only used to evaluate the quality of our results, as all the performed benchmarking is unsupervised.

For scRNA-seq data four datasets have been considered. First, the scRNA-seq data (called “Liu scRNA”) present in the scCAT-seq joint profiling of Liu et al. [24] containing 206 cells profiled from three cancer cell lines (HCT116, HeLa-S3, K562). Next, a bigger dataset composed of 561 cells profiled from seven cell lines (A549, GM12878, H1437, HCT116, IMR90, H1, K562) from Li et al. [25] (called “Li cell lines”). Finally, two colorectal cancer (CRC) datasets, corresponding to primary CRC tumors and matched normal mucosa have been also taken into account. The first (called “Li Tumor”) contains 364 cells clustered into seven cell types: B cells, endothelial cells, epithelial cells, fibroblasts, macrophages, mast cells and T cells. The second (called “Li NM”) is composed of 266 cells clustered according to the same seven cell types.

Other single-cell omics are also included in our analysis: methylation and scATAC-seq data. For scATAC-seq data we considered: i) the dataset included into the scCAT-seq joint profiling of Liu et al. [24] (called “Liu scATACseq”) composed of 206 cells extracted from three cancer cell lines (HCT116, HeLa-S3, K562); ii) the leukemia scATAC-seq data from Corces et al. [26] (called “Leukemia scATAC”), containing 391 cells and composed of monocytes and lymphoid-primed multipotent progenitors (LMPP) isolated from a healthy human donor, together with leukemia stem cells (SU070_LSC, SU353_LSC) and blast cells (SU070_Leuk, SU353_Blast), isolated from two patients with acute myeloid leukemia. To represent single-cell DNA methylation we considered two neuronal snmC-seq datasets from Luo et al. [27]. The first (called “scMethylation mouse”) is composed of 3,377 cells extracted from mouse frontal cortical neurons clustered into 16 neuronal subtypes. The second (called “scMethylation human”) is composed of 2,740 cells, extracted from human frontal cortical neurons, and clustered into 21 neuronal subtypes.

A summary of the considered datasets is available in Table 1. The downloaded datasets had already undergone standard preliminary preprocessing and, following standard practices [7], we log-transformed the scRNA-seq counts and selected the 10,000 most varying features.

### Baseline metrics

We consider a single-cell omics dataset as a matrix *X*, whose columns correspond to cells, and whose rows correspond to features (e.g. peaks, genes). Given two cells indexed by *l* and *m* as columns of *X*, different metrics are classically used to infer the similarity between their omic profiles *x* = *X*[, *l*] = (*x*_1_… *x*_*n*_) and *y* = *X*[ , *m*] = (*y*_1_… *y*_*n*_). We here focus on four state-of-the-art metrics, called in the following *baseline metrics*, and defined as:

i. Euclidean distance (L2): 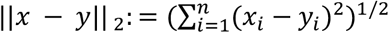
ii. Manhattan distance (L1): 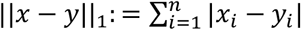
iii. Cosine similarity: 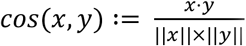, where ∥ · ∥ is the euclidean norm
iv. Pearson correlation: 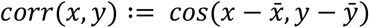, where 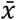 and 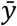 are the mean of the values of*x*and *y*, respectively.

For cosine similarity and Pearson correlation we used the distance-like formulation 1 − *cos*(*x*, *y*) and 1 − *corr*(*x*, *y*). Of note, these are not strictly speaking distances, in particular they do not respect the triangular inequality. The baseline metrics have been computed using functions from the Python package SciPy (`scipy.spatial.distance`) : `euclidean, cityblock, cosine, correlation`.

Baseline metrics have been computed on the input data, after the data preprocessing detailed in the section above. Alternative optional normalizations of the data have been also considered (Figure 1), but they resulted to be less performant (Supplementary Table 2).

**Figure 1:**
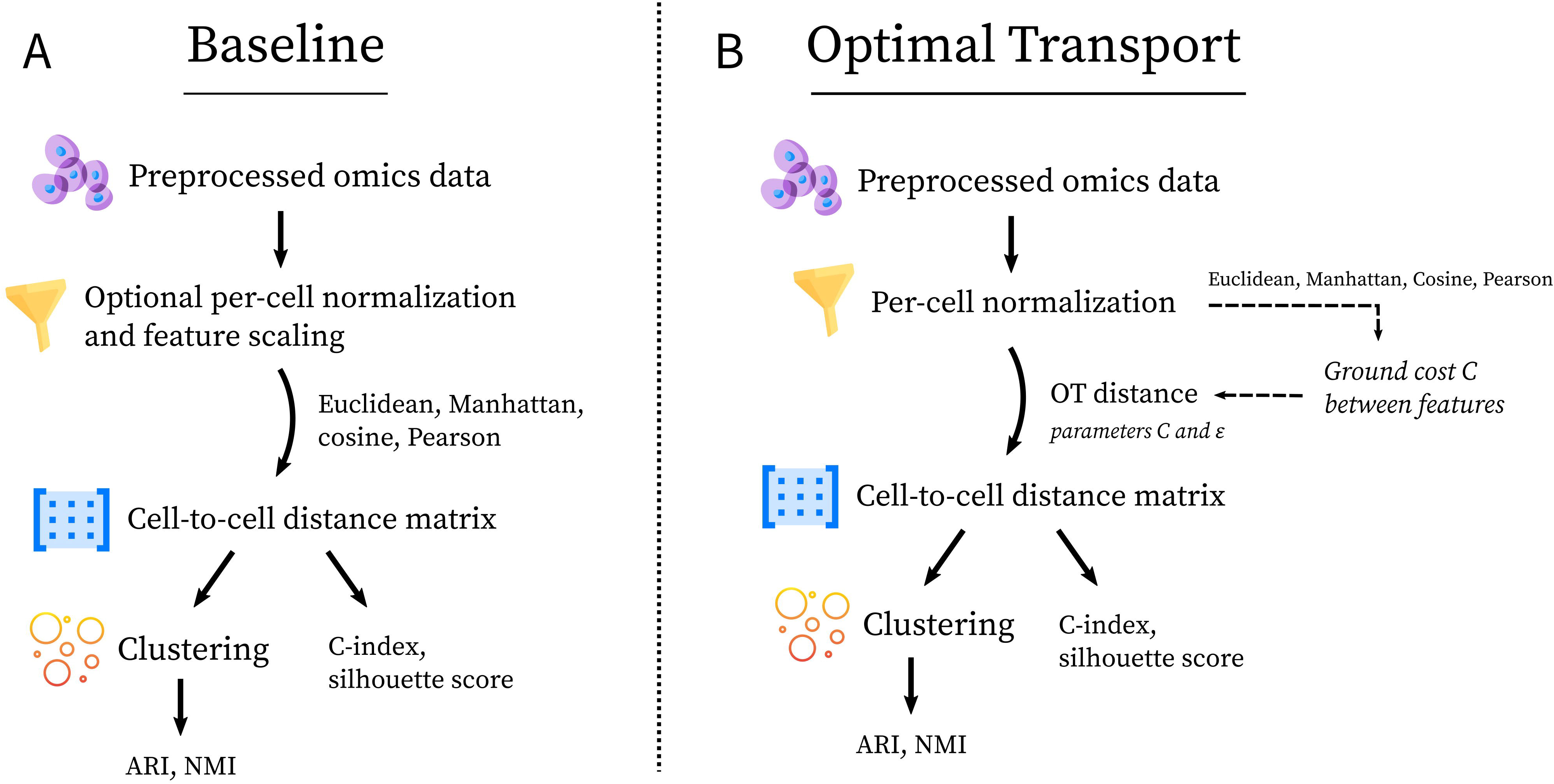
Workflow for metrics comparison. The employed procedure, from the input preprocessed data to the performance evaluation is summarized for (A) baseline metrics and (B) Optimal Transport, respectively. The graphic contents in the figure are taken from flaticon.com.

### Optimal Transport distance

Optimal Transport (OT), as defined by [28,29], aims at finding a coupling between two probability distributions that minimizes transportation cost. The classical OT distance, also known as the Wasserstein distance, between two distributions *a* = (*a*_1_… *a*_*n*_) and *b* = (*b*_1_… *b*_*n*_) is defined as the minimal cost of transportation to morph *a* into *b*. Given *a* and *b* discrete histograms, their OT distance is thus defined as:

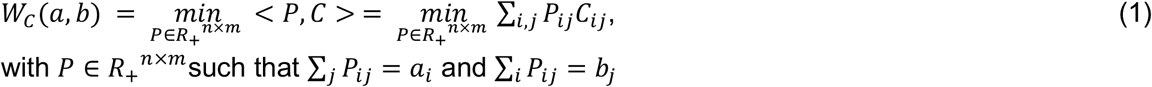

where *P*is the coupling. According to *P*, the mass in the histogram *a* is thus moved from one bin to another one in order to transform *a* into *b*. *C*is called *ground cost* and it encodes the penalty for moving a unit of mass from one bin to another one. Hence *C* should be chosen in such a way that similar bins *i* and *j* have a low cost *C*_*ij*_.

We propose the use of the OT distance to capture cell-cell similarity in different single-cell omics data. For simplicity, let us consider a scRNA-seq dataset (*X*); the same concepts apply to other single-cell omics. Given a pair of cells *l* and *m*, as done for baseline metrics, we consider their expression profiles, corresponding to the vectors *x* = *X*[, *l*] = (*x*_1_… *x*_*n*_) and *y* = *X*[, *m*] = (*y*_1_… *y*_*n*_). Given that Eq (1) is defined only for histograms, we transform *x*and *y* into two histograms 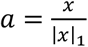 and 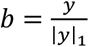. In the following, we refer to such a transformation as *per-cell normalization*. After transformation, we can apply Eq (1) and compute the OT distance *W*_*C*_(*a*, *b*), between *a* and *b*, which corresponds to evaluating the minimal cost required to transform the gene expression histogram (*a*) of the first cell into the gene expression histogram (*b*) of the second cell. Based on Eq. (1), OT computes the distance between a pair of cells (*a*, *b*) by taking into account the joint gene expression activity present in the two cells. For this reason, it is expected to better capture the similarity between a pair of cells having activated similar transcriptional programs.

As discussed above, the OT distance is parametrized by the ground cost *C*. The choice of the ground cost plays a central role in the final performances of OT and, in our case, there is no straightforward choice. Hence, for all couples of genes *i* and *j* in the single-cell matrix *X*, we define:

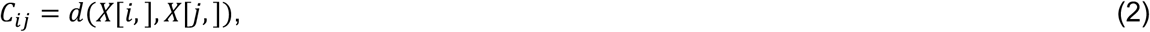

where *d* corresponds to a metric among Euclidean and Manhattan distance, Cosine similarity and Pearson correlation. Intuitively, *C*_*ij*_ thus reflects the cost of moving a unit of gene expression from gene *i* to gene *j*.

Given that the number of features (e.g. genes, peaks) of single-cell data is in the order of tens of thousands, the classical Optimal Transport problem would be computationally intractable. Indeed, solving Eq. (1) relies on costly linear programming methods [14]. Hence, we considered the entropic regularization of the classical OT distance, also called Sinkhorn divergence [21,30]. The entropic-regularized OT distance between two distributions *a* and *b* is defined as:

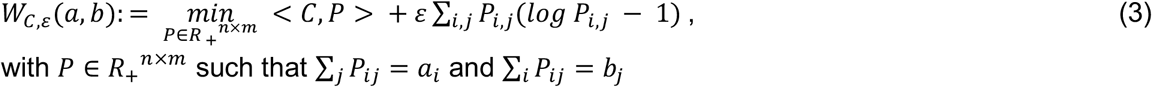

The first term of Eq. (3) corresponds exactly to Eq. (1) with *P* coupling and *C* ground cost. The additional term corresponds to the entropic regularization. Therefore, if the regularization parameter *ε* is set to zero, Eq (3) corresponds exactly to Eq (1) and classical OT is obtained. Increasing values of *ε* correspond to a more diffused coupling and a faster execution of the algorithm. The parameter *ε* thus plays a central role in the final performances of Sinkhorn divergence and should be carefully chosen. The advantage of the formulation at Eq (3) is that *W*_*C,ε*_can be efficiently computed on a GPU, thereby coping with the high dimensionality of single-cell data. Not only is entropy pivotal to scale the algorithm, but it is also important to break the curse of dimensionality which makes classical OT distance extremely hard to estimate from high-dimensional single-cell data. This phenomenon, analyzed theoretically in [30] is supported by our analysis (Supplementary Table 1).

An issue with Eq (3) for *ε* > 0is that, given two cells having the same expression distribution (*a* = *b*), *W*_*C,ε*_(*a*, *b*) > 0. We thus used the debiased Sinkhorn divergence [31] in order to ensure a distance equal to zero for identical cells.

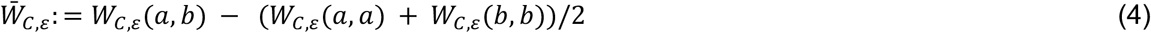

For sake of simplicity in the rest of the paper, we will use the term *OT distance* to refer to the debiased Sinkhorn divergence.

As discussed above, the OT distance depends on two main parameters: the regularization parameter *ε*and the ground cost *C*. For every dataset, we performed a grid search varying *ε* among 0.1, 0.05, 0.01, and the ground cost among Euclidean, Manhattan, Cosine and Pearson correlation. As shown in Supplementary Table 1, for all datasets, the best performances were obtained for the regularization parameter *ε* set at 0.1. In contrast, the best performing ground cost *C*varied depending on the analyzed data. In scRNA-seq simulated data and in single-cell DNA methylation data, Pearson correlation achieved the best performances, while on scRNA-seq and scATAC-seq data, cosine similarity performed the best. The performances presented in the Results section thus correspond to this choice of *ϵ*and *C*. The computation of the OT distance has been implemented using the Python package PyTorch and run on a GPU.

### Performance evaluation

For all single-cell datasets, simulated and real, ground-truth labels are available. For the real single-cell omics, the ground-truth labels correspond to cell types, defined through clustering in the original publication, or to the cell lines from which the cells have been extracted.

We first use the C-index and Silhouette score to evaluate to which extent the various metrics detect the similarity between cells associated with the same label, as well as the difference between cells with different labels (Figure 1). The C-index [32] is an internal clustering evaluation index. Given a cell-to-cell distance matrix, the C-index measures if the closest pairwise distances correspond to cells belonging to the same cluster. It is defined as:

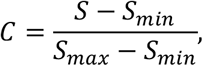

where *n* is the number of intra-cluster pairwise distances, *S*_*min*_is the sum of the *n* smallest distances if all pairs of cells are considered, *S*_*max*_ is the sum of the *n* largest distances out of all pairs and *S* is the sum of distances over all pairs of cells form the same cluster.

Of note, the C-index is always in the interval [0, 1] and it should be minimum in the case of a perfect clustering. To make the results easily readable, we consider 1 − *C*, so that the best performances are obtained by maximizing the score. Concerning the implementation, we used our own Python implementation of the C-index.

As a complementary evaluation, we further considered the Silhouette score, defined as:

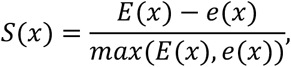

where *E*(*x*) is the average distance between the cell *x* and the other cells of the same cluster and *e*(*x*) is the average distance between *x* and cells in the closest different cluster. The distance used to compute *E*(*x*) and *e*(*x*) varies among the benchmarked metrics (OT, Euclidean, Manhattan, Cosine similarity and Pearson correlation). The global Silhouette score is then obtained by averaging *S*(*x*)over all cells. We used the Silhouette score implementation of the Python package scikit-learn (`sklearn`) [33].

To further test the quality of the inferred cell-to-cell distance matrices, we used them as inputs for a clustering algorithm and assessed the agreement between the inferred clusters and the ground-truth labels (Figure 1). We only considered clustering algorithms for which applications to single-cell data have been already proposed and directly applicable to the cell-to-cell distance matrices, without performing any post-processing [8]. We thus selected hierarchical clustering, with complete linkage, and spectral clustering [34,35], both implemented in scikit-learn. Regarding hierarchical clustering, we chose complete linkage in place of average linkage, used in other single-cell clustering works [11], because this approach provided better performances for both baseline and OT distances (Supplementary Table 3). Concerning spectral clustering, since it requires in input an affinity matrix, we converted the inferred distance matrices *D* to affinity matrices *A* = 1 − *D*, with *D* normalized such that the maximum value is set to 1, and run the clustering algorithm with default parameters. Both clustering algorithms require the specification of the number of clusters in which the cells should be partitioned. We chose to set the number of clusters in an unsupervised way, by optimizing the Silhouette score [36]. For each distance matrix, we thus varied the number of clusters (*k*) in the range 3-25, and chose the *k* maximising the Silhouette score of the clustering. Thereby, we can test how frequently the number of ground-truth labels present in the data are captured by the different metrics. Of note, the overall behaviour observed when the number of clusters is optimized is in good agreement with the results obtained by fixing the number of clusters to the number of ground-truth labels (Supplementary Table 3). To evaluate the quality of the obtained clusters, we used Adjusted Rand Index (ARI) and Normalized Mutual Information (NMI) (Figure 1).

Given *U*, clustering inferred from a distance matrix and *V*, ground-truth labels, the ARI is defined as:

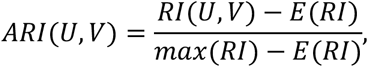

where *RI*(*U*, *V*) is the Rand index, i.e. the fraction of pairs of samples that are either in the same group or in different groups in both *U* and *V*, *E*(*RI*)is the expected Rand index between *U* and a random *V*, and *max*(*RI*) is the largest possible Rand index between *U* and any *V*.

To consider a complementary score, we also used the NMI, defined as:

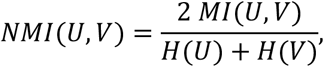

where *MI*(*U*, *V*)is the mutual information between *U* and *V*,i.e. *MI*(*U*, *V*) = *H*(*U*) − *H*(*U*|*V*) and *H*(.) denotes entropy. To compute ARI and NMI, we used the corresponding scikit-learn implementations.

## Results

Debiased entropic-regularized Optimal Transport (OT) distance (see Methods), called in the following OT distance, is here proposed as a metric to infer cell-cell similarity across different single-cell omics data. The performances of the OT distance are then benchmarked with respect to state-of-the-art metrics, called in the following baseline metrics, namely the Euclidean and Manhattan distances, Cosine similarity and Pearson correlation.

The benchmark is performed in three main contexts (Table 1). First, simulated scRNA-seq data are considered. Data composed of different numbers of cells are generated to test whether the metrics scale also to high-dimensional data, as the currently available single-cell data, and if the number of cells impacts performances. Then the overlap and size of the clusters underlying the simulated scRNA-seq data are also varied to challenge the various metrics in detecting less clear and rare groupings of cells. Second, four real scRNA-seq data, profiled from colorectal cancer (CRC) and cell lines, are considered. Finally, we further challenged the distance measures on other single-cell omics: scATAC-seq and single-cell DNA methylation. See Methods and Table 1 for details concerning the data.

In all three contexts, ground-truth labels were available for all cells. In simulated scRNA-seq data, the ground-truth labels have been imposed during the data simulation (see Methods). In real single-cell omics, the ground-truth labels are extracted from the original publications. For the four profilings of cell lines (Liu scRNA, Li cell lines, Liu scATAC and Leukemia scATAC), labels correspond to the cell line of origin and thus reflect strong transcriptional/epigenetic differences. In contrast, in the case of the neuronal single-cell DNA methylation (scMethylation mouse and scMethylation human) and scRNA-seq from colorectal cancer samples (Li Tumor and Li NM), the ground-truth labels reflect clusters previously identified in the data, based on the activity of predefined markers. These last applications are clearly more challenging, as much weaker differences exist between different cell types or states.

As summarized in Figure 1, we first tested with C-index and Silhouette score whether the various metrics are able to detect the similarity between cells belonging to the same group. We then applied different unsupervised clustering algorithms to the computed distance matrices and tested the quality of the resulting clusters using the Adjusted Rand Index (ARI) and Normalized Mutual Information (NMI).

### Comparing Optimal Transport against baseline metrics based on cell-cell similarity detection

Figure 2 summarizes the results of the comparison between Optimal Transport (OT) and baseline metrics. Pearson correlation outperformed the other baseline metrics on all data types. We thus used it as representative of baseline metrics in Figure 2 and in the following sections. The results of alternative baseline metrics are available in Supplementary Table 2. Of note, better performances of Pearson correlation with respect to other state-of-the-art metrics had been previously observed [12].

**Figure 2.**
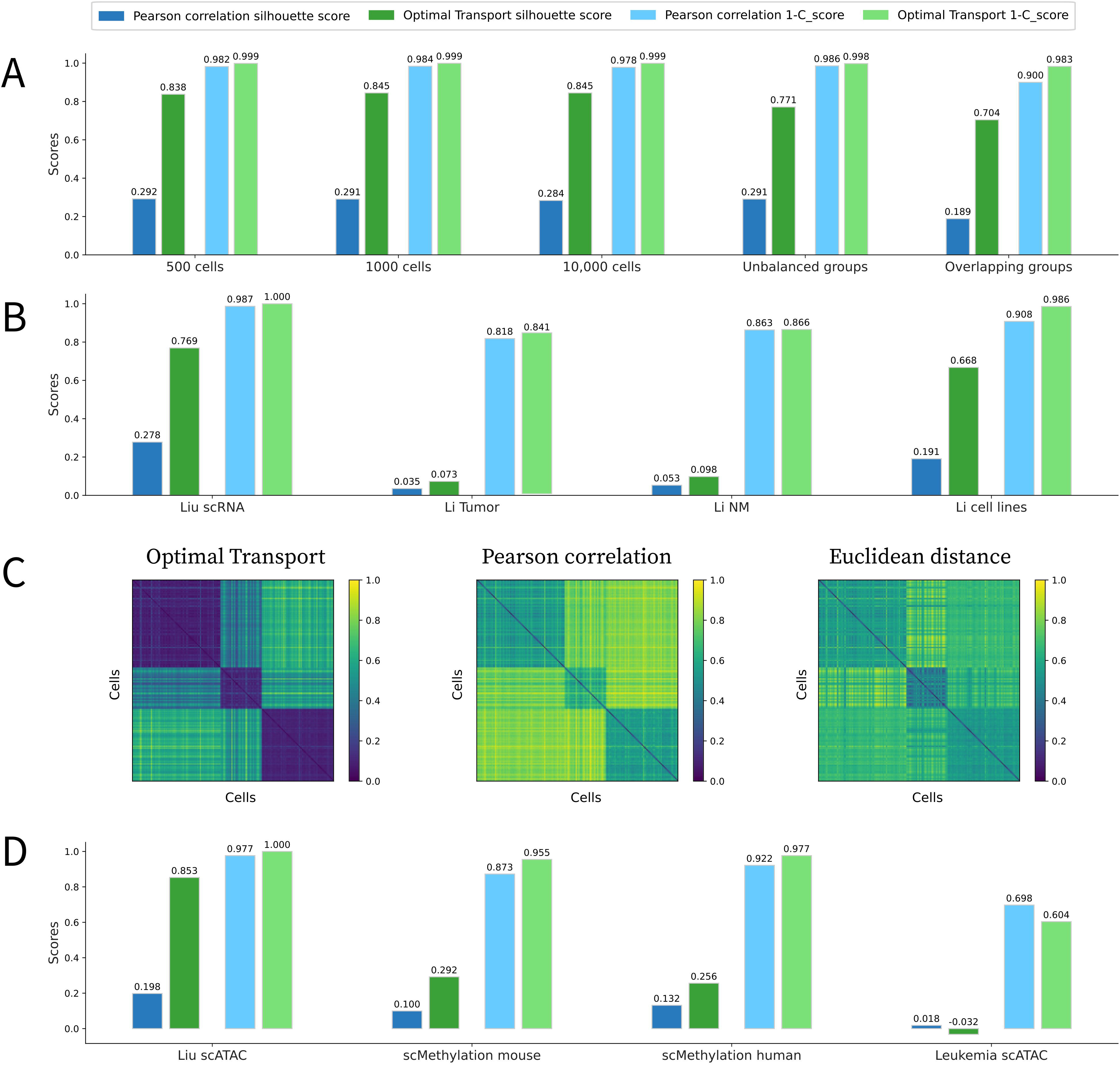
Comparison of Optimal Transport (OT) against Pearson correlation in cell-cell similarity inference. Barplots for C-index and Silhouette score are reported for (A) simulated scRNA-seq data composed of 500, 1,000 and 10,000 cells, with unbalanced groups and overlapping clusters; (B) four scRNA-seq datasets; (D) two single-cell DNA methylation and two scATAC-seq data. Examples of the distance matrices obtained with OT, Pearson correlation and Euclidean distance in Liu scRNA-seq are reported in (C).

In all simulated data, OT outperforms all baseline metrics (Figure 2A), both in terms of C-index and Silhouette score. In particular, the results of OT are not impacted by the number of cells, given that it shows a consistently performant behavior for 500, 1,000 and 10,000 cells. These results also suggest that OT is scalable to high-dimensional datasets, a crucial feature for the analysis of single-cell data. Finally, OT achieves superior performance with highly unbalanced clusters, which is more realistic as biological samples are often composed of a mixture of rare and common populations of cells, and also with overlapping clusters, which reflects the scenario of subpopulations of cells sharing similar transcriptional patterns, as cell populations tracked over different developmental phases.

We then considered four scRNA-seq datasets: two of them correspond to cancer cell lines [24,25], while the remaining two correspond to colorectal tumour tissue and matched normal mucosa [25]. All metrics tend to perform better in cancer cell lines than CRC samples. This result is most probably the consequence of the stronger transcriptional difference existing between cell lines. Overall, in all the four scRNA-seq datasets, OT outperformed baseline metrics (Figure 2B). The improvement provided by OT in cell lines is important, especially according to Silhouette score (+0.5 Silhouette score). To show to which extent C-index and Silhouette score reflect a clear clustering structure in the cell-to-cell distance matrices, we focused on the smallest dataset, Liu scRNA [24]. Figure 2C reports the cell-to-cell distance matrices obtained for this dataset with OT distance, Pearson correlation and Euclidean distance. Cells in the matrices are sorted based on their cell line of origin. OT powerfully detected the similarity between cells belonging to the same cancer cell line, showing three clear blocks of cells at distance close to zero. In contrast, the blocks corresponding to the three cell lines are less marked with Pearson correlation. Finally, with Euclidean distance, the values outside the three blocks tend to be less close to one, indicating a less clear separation between cells belonging to different cell lines.

Finally, we challenged the various metrics on other single-cell omics data (Figure 2D). The results obtained with Pearson correlation and OT are comparable. First, in all datasets, except the scATAC-seq cell lines, the Silhouette scores were quite low, indicating that detecting cell-cell similarities in scATAC-seq and single-cell DNA methylation data is more challenging. In addition, OT performed better for the Liu scATAC-seq and methylation data (mouse and human), while Pearson correlation performed better for the leukemia scATAC-seq dataset.

Overall, OT clearly outperformed existing metrics in the detection of cell-cell similarities in simulated and real scRNA-seq and single-cell DNA methylation data. The results are less clear for scATAC-seq data. See Supplementary Table 2 for the performances of other state-of-the-art baseline metrics.

### Comparing Optimal Transport against baseline metrics based on hierarchical clustering

Hierarchical clustering was applied to the cell-to-cell distance matrices computed with OT and Pearson correlation. In all datasets, the number of clusters was optimized based on the Silhouette score (Methods). Table 1 summarizes the number of clusters obtained across all datasets and compares them with those defined in the original publications. The resulting clusters were then compared with the ground-truth labels based on Adjusted Rand Index (ARI) and Normalized Mutual Information (NMI) (Figure 3).

In simulated data, all the five datasets contain three underlying clusters. As described in Table 1, in four out of five cases, both OT and Pearson correlation identified the correct number of clusters. The only exception is represented by the dataset composed of 10,000 cells, where OT correctly detects the presence of three clusters, while Pearson correlation overfitted the data, by subdividing the three groups into five clusters. According to ARI and NMI, both Pearson correlation and OT give rise to perfect results (Figure 3A). Pearson correlation also led to optimal ARI and NMI scores in the dataset of 10,000 cells, as overfitting is not captured by these scores.

**Figure 3.**
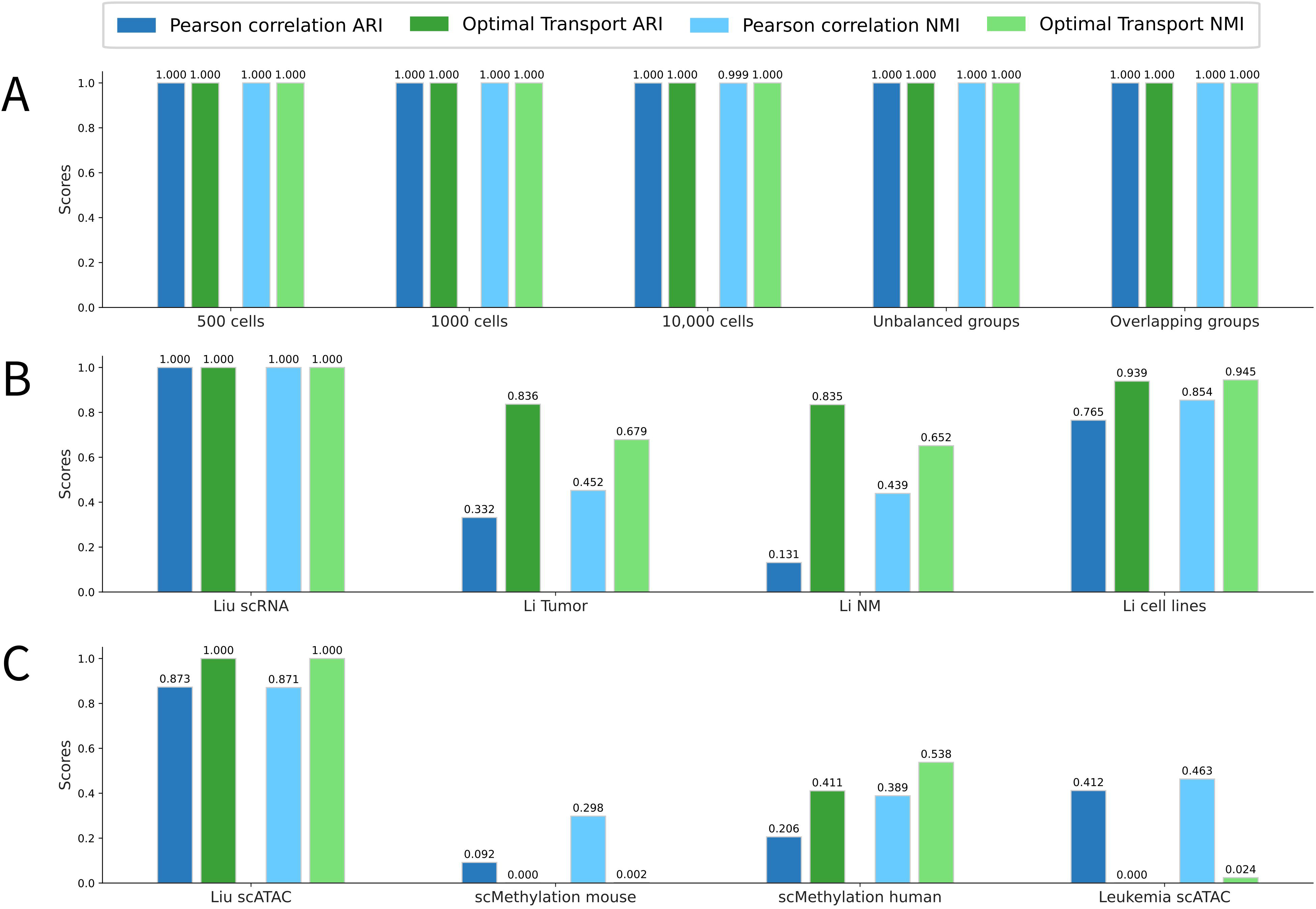
Comparison of Optimal Transport (OT) against Pearson correlation in hierarchical clustering. Barplots for Adjusted Rand Index (ARI) and Normalized Mutual Information (NMI) are reported for (A) simulated scRNA-seq data composed of 500, 1,000 and 10,000 cells, with unbalanced groups and overlapping clusters; (B) four scRNA-seq datasets C. two single-cell DNA methylation and two scATAC-seq data.

Turning to real scRNA-seq data, in the case of Liu scRNA data, both Pearson correlation and OT detected the correct number of clusters (Table 1). Concerning the two CRC datasets, for Li Tumor, containing seven cell types, both Pearson and OT predicted the presence of six clusters, thus missing the rare population of mast cells, only represented by one cell. In contrast, for Li NM, Pearson correlation dramatically overfitted the data and inferred 23 clusters, while OT suggested the presence of five clusters. Finally, for Li cell lines, composed of seven cell lines, OT suggested the presence of eight clusters, while Pearson identified ten. Overall, in all scRNA-seq datasets, excepting Liu [24], no distance captured exactly the numbers of clusters reported in the corresponding publications. However, the number of clusters inferred by OT was always much closer to the ground-truth, while Pearson correlation tended to highly overfit the data. Regarding the NMI and ARI scores, in three out of four datasets, OT outperformed Pearson correlation (Figure 3B). The only exception is represented by Liu scRNA, for which both metrics achieved perfect clustering. Interestingly the clustering improvement produced by OT was stronger for the two CRC data, which are also the most challenging to cluster.

In other single-cell omics data, the results are less clear and, as observed previously, the performances of OT are comparable with those of the best performing baseline metrics (Pearson correlation). Regarding the numbers of clusters, in Liu scATAC, both Pearson correlation and OT correctly detected the presence of three clusters. In Leukemia scATAC, composed of six cell lines, OT and Pearson correlation predicted three and 25 clusters, respectively. Finally, in methylation data, both OT and Pearson underestimated the real number of clusters. As shown in Figure 3C, the ARI and NMI values follow the same trends as the C-index and Silhouette score shown in Figure 2D. OT showed better performances in cell lines scATAC and human methylation, while Pearson correlation performed better for mouse methylation and Leukemia data. Of note, Pearson correlation performed well according to ARI and NMI for Leukemia data, but this performance is obtained considering 25 clusters instead of the five real clusters presumably present in the data. This is a practical demonstration of the interest of inferring the optimal number of clusters based on the Silhouette score, rather than fixing it to the value reported in their original publication.

The improvement provided by OT for cell-cell similarity inference (Figure 2) has an impact on clustering performances. Of note, these results are also in agreement with those observed when the number of clusters is fixed (Supplementary Table 3).

### Comparing Optimal Transport against baseline metrics based on spectral clustering

To test whether the improvement provided by OT in cell clustering depends on the clustering algorithm, we repeated the same tests with spectral clustering. Also in this case, the number of clusters was optimised based on the Silhouette score (Methods). The numbers of resulting clusters are listed in Table 1, while the corresponding values of ARI and NMI are provided in Figure 4.

**Figure 4.**
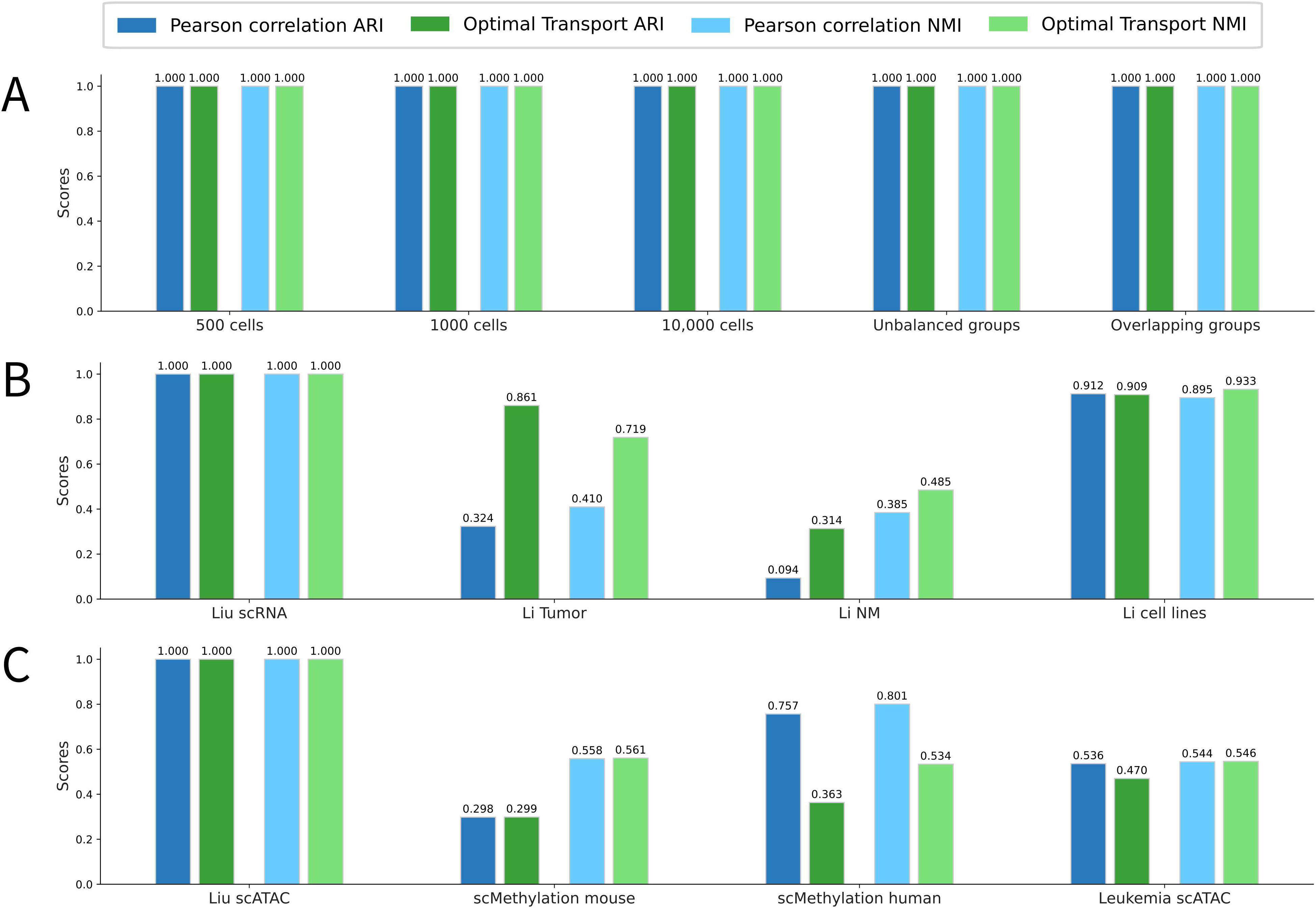
Comparison of Optimal Transport (OT) against Pearson correlation in spectral clustering. Barplots for Adjusted Rand Index (ARI) and Normalized Mutual Information (NMI) are reported for (A) simulated scRNA-seq data composed of 500, 1,000 and 10,000 cells, with unbalanced groups and overlapping clusters; (B) four scRNA-seq datasets: (C) two single-cell DNA methylation and two scATAC-seq data.

In simulated data, Pearson correlation and OT metrics always detected the correct number of clusters (Table1) and achieved perfect clustering according to ARI and NMI (Figure 4A).

For scRNA-seq data, both Pearson correlation and OT correctly predicted the clusters present in Liu scRNA. The two CRC datasets contain seven cell types, corresponding to our ground-truth labels. For Li Tumor, both Pearson and OT predicted the presence of three clusters. In contrast, for Li NM, Pearson correlation overfitted the data and inferred 13 clusters, while OT suggested the presence of five clusters. Therefore, in colorectal cancer, a better approximation of the true clustering present in the data has been obtained with hierarchical clustering. Finally, for Li cell lines, OT suggested the presence of five clusters, while Pearson correctly identified seven clusters. Concerning the NMI and ARI scores, OT outperformed Pearson correlation for three of the four datasets (Figure 4B). The only exception is represented by Liu scRNA data, where both Pearson and OT achieved perfect clustering.

Concerning other single-cell omics, Pearson correlation and OT correctly identified three clusters for Liu scATAC data. Regarding Leukemia scATAC data, involving six cell lines, both OT and Pearson overfitted the data, predicting 18 and 14 clusters, respectively. Finally, in methylation data both OT and Pearson drastically underestimated the real number of clusters. Regarding ARI and NMI (Figure 4C), for Liu ATAC data, the results are optimal with both metrics. In all other datasets, performances are slightly changed compared to those observed with hierarchical clustering (Figure 3C). For mouse scMethylation and Leukemia scATAC data, Pearson and OT showed comparable performances, while for human scMethylation data, Pearson correlation performed better.

Overall, the clustering conclusions previously derived with hierarchical clustering are confirmed with spectral clustering. Indeed, there is a good agreement between spectral clustering and hierarchical clustering for simulated data and scRNA-seq data. Only marginal variations were observed, such as a reduced difference between Pearson and OT for colorectal cancer data, when spectral clustering is employed. In other single-cell omics data, the results of the two clustering algorithms are less concordant, but the overall performances of Pearson and OT remain comparable. Of note, we obtained similar results for fixed numbers of clusters (Supplementary Table 3).

### Open-source implementation and distribution

To foster the reproducibility of all the results presented in this work, we provide a Jupyter notebook covering all the analyses performed. Coded in Python, this notebook is available at https://github.com/ComputationalSystemsBiology/ot-scOmics, together with all the preprocessed single-cell data used in this study. Since computing Optimal Transport distances is computationally intensive, the code is designed to be run on a GPU, taking advantage of the PyTorch library. For users who do not have access to a GPU, extensive explanations are provided to run the Jupyter notebooks on the Google Collaboratory platform (http://colab.research.google.com/).

## Discussion

In this study, we assessed the potential of Optimal Transport (OT) to infer cell-cell similarities from single-cell omics data. We extensively benchmarked OT performances against state-of-the-art metrics. Interestingly, OT outperformed alternative metrics in capturing cell-cell similarities in simulated and real scRNAseq data. The biological relevance of this improvement was assessed by performing cell clustering. In all cases, the use of OT distance resulted in improved clustering results. Of note, two clustering algorithms have been used to test whether the observed performances were affected by the choice of the algorithm. We further challenged the metrics to detect cell-cell similarity in other single-cell omics: scATAC-seq and single-cell DNA methylation data. In these cases, OT showed performances comparable with the best performing baseline metric, namely Pearson correlation. Overall, the results obtained for scATAC-seq and single-cell DNA methylation data are less convincing than those for scRNA-seq data. These results are presumably due to the additional challenges introduced by these omics data, in particular their high sparsity and the close-to-binary nature of scATAC-seq data [9,37].

Our evaluation of metrics performances is dependent on the ground-truth labels associated with the input datasets. In cell lines, the ground-truth is well established and wide transcriptional differences exist between different cell lines. Biological samples instead consist of cell types and states having less pronounced transcriptional differences. What is a ground-truth in this case is thus less clear. We here used as ground-truth the clusters identified in the original publication of each dataset. However, such labels could be improved. In addition, given that the original publications employ Euclidean distance or Pearson correlation to define the labels, the usage of such labels as ground-truth is expected to advantage these metrics compared to alternative ones.

Of note, Dimensionality Reduction (DR) is frequently applied to reduce the noise in single-cell data before cell clustering and further downstream analyses. Here, we applied feature selection without DR to test to which extent different metrics are affected by such noise. In addition, the most popular DR approaches, such as PCA, diffusion maps, NMF and ICA, rely on Euclidean geometry. Hence, we avoided using DR to not favor the Euclidean metrics in our benchmarking. However, the good performances provided by OT suggest that further efforts should be devoted to design DR methods for single-cell data based on other metrics.

Finally, we applied OT to different single-cell omics in isolation. However, different omics data presumably provide complementary information on individual cellular states. Combining different single-cell omics with appropriate metrics thus represent a critical challenge in computational biology. Our results suggest that OT could be a valuable metric for the integration of different omics data.

## Data availability

The datasets used in the study are all public and accessible through their corresponding publication (Table 1). Preprocessed data together with the code to reproduce the analyses are available at https://github.com/ComputationalSystemsBiology/OT-scOmics.

## Acknowledgements

We thank Denis Thieffry for the scientific feedback on the work and Jean-Philippe Vert for the insightful discussion during the design of this project.

## References

1. Stegle O, Teichmann SA, Marioni JC. Computational and analytical challenges in single-cell transcriptomics. Nat. Rev. Genet. 2015; 16:133–145

2. Rajewsky N, Almouzni G, Gorski SA, et al. LifeTime and improving European healthcare through cell-based interceptive medicine. Nature 2020; 587:377–386

3. Papalexi E, Satija R. Single-cell RNA sequencing to explore immune cell heterogeneity. Nat. Rev. Immunol. 2018; 18:35–45

4. Potter SS. Single-cell RNA sequencing for the study of development, physiology and disease. Nat. Rev. Nephrol. 2018; 14:479–492

5. Lee J, Hyeon DY, Hwang D. Single-cell multiomics: technologies and data analysis methods. Exp. Mol. Med. 2020; 52:1428–1442

6. Ma A, McDermaid A, Xu J, et al. Integrative Methods and Practical Challenges for Single-Cell Multi-omics. Trends Biotechnol. 2020; 38:1007–1022

7. Luecken MD, Theis FJ. Current best practices in single-cell RNA-seq analysis: a tutorial. Mol. Syst. Biol. 2019; 15:e8746

8. Kiselev VY, Andrews TS, Hemberg M. Challenges in unsupervised clustering of single-cell RNA-seq data. Nat. Rev. Genet. 2019; 20:273–282

9. Xiong L, Xu K, Tian K, et al. SCALE method for single-cell ATAC-seq analysis via latent feature extraction. Nat. Commun. 2019; 10:4576

10. Satija R, Farrell JA, Gennert D, et al. Spatial reconstruction of single-cell gene expression data. Nat. Biotechnol. 2015; 33:495–502

11. Guo M, Wang H, Potter SS, et al. SINCERA: A Pipeline for Single-Cell RNA-Seq Profiling Analysis. PLoS Comput. Biol. 2015; 11:e1004575

12. Kim T, Chen IR, Lin Y, et al. Impact of similarity metrics on single-cell RNA-seq data clustering. Brief. Bioinform. 2019; 20:2316–2326

13. Macosko EZ, Basu A, Satija R, et al. Highly Parallel Genome-wide Expression Profiling of Individual Cells Using Nanoliter Droplets. Cell 2015; 161:1202–1214

14. Peyré G, Cuturi M. Computational optimal transport: With applications to data science. Found. Trends® Mach. Learn. 2019; 11:355–607

15. Santambrogio F. Optimal transport for applied mathematicians. Birkäuser NY 2015; 55:94

16. Schiebinger G, Shu J, Tabaka M, et al. Optimal-Transport Analysis of Single-Cell Gene Expression Identifies Developmental Trajectories in Reprogramming. Cell 2019; 176:928–943.e22

17. Demetci P, Santorella R, Sandstede B, et al. Gromov-Wasserstein optimal transport to align single-cell multi-omics data. BioRxiv 2020;

18. Cao K, Hong Y, Wan L. Manifold alignment for heterogeneous single-cell multi-omics data integration using Pamona. bioRxiv 2020;

19. Bellazzi R, Codegoni A, Gualandi S, et al. The Gene Mover’s Distance: Single-cell similarity via Optimal Transport. ArXiv Prepr. ArXiv210201218 2021;

20. Huizing G-J, Cantini L, Peyré G. Unsupervised Ground Metric Learning using Wasserstein Eigenvectors. ArXiv Prepr. ArXiv210206278 2021;

21. Cuturi M. Sinkhorn distances: lightspeed computation of optimal transport. NIPS 2013; 2:4

22. Regev A, Teichmann SA, Lander ES, et al. The Human Cell Atlas. eLife 2017; 6:

23. Zappia L, Phipson B, Oshlack A. Splatter: simulation of single-cell RNA sequencing data. Genome Biol. 2017; 18:174

24. Liu L, Liu C, Quintero A, et al. Deconvolution of single-cell multi-omics layers reveals regulatory heterogeneity. Nat. Commun. 2019; 10:470

25. Li H, Courtois ET, Sengupta D, et al. Reference component analysis of single-cell transcriptomes elucidates cellular heterogeneity in human colorectal tumors. Nat. Genet. 2017; 49:708–718

26. Corces MR, Buenrostro JD, Wu B, et al. Lineage-specific and single-cell chromatin accessibility charts human hematopoiesis and leukemia evolution. Nat. Genet. 2016; 48:1193–1203

27. Luo C, Keown CL, Kurihara L, et al. Single-cell methylomes identify neuronal subtypes and regulatory elements in mammalian cortex. Science 2017; 357:600–604

28. Monge G. Mémoire sur la théorie des déblais et des remblais. 1781;

29. Kantorovich L. On the transfer of masses (in Russian). Dokl. Akad. Nauk 1942; 37:227–229

30. Genevay A, Chizat L, Bach F, et al. Sample complexity of sinkhorn divergences. 22nd Int. Conf. Artif. Intell. Stat. 2019; 1574–1583

31. Feydy J, Séjourné T, Vialard F-X, et al. Interpolating between optimal transport and MMD using Sinkhorn divergences. 22nd Int. Conf. Artif. Intell. Stat. 2019; 2681–2690

32. Hubert L, Schultz J. Quadratic assignment as a general data analysis strategy. Br. J. Math. Stat. Psychol. 1976; 29:190–241

33. Pedregosa F, Varoquaux G, Gramfort A, et al. Scikit-learn: Machine learning in Python. J. Mach. Learn. Res. 2011; 12:2825–2830

34. Von Luxburg U. A tutorial on spectral clustering. Stat. Comput. 2007; 17:395–416

35. Zheng R, Li M, Liang Z, et al. SinNLRR: a robust subspace clustering method for cell type detection by non-negative and low-rank representation. Bioinformatics 2019; 35:3642–3650

36. Rousseeuw PJ. Silhouettes: a graphical aid to the interpretation and validation of cluster analysis. J. Comput. Appl. Math. 1987; 20:53–65

37. P E de Souza C, Andronescu M, Masud T, et al. Epiclomal: Probabilistic clustering of sparse single-cell DNA methylation data. PLoS Comput. Biol. 2020; 16:e1008270

