## Supplementary Text for "Optimal Transport improves cell-cell similarity inference in single-cell omics data"

Parameters were estimated from the Liu et al. scRNA data through the function `splatEstimate`.
The following parameters were refined manually:

- `nGenes` (number of genes in simulated dataset): 5000 for all simulations
- `batchCells` (number of cells in simulated dataset): 500, 1000 or 10000 depending on the simulation.
- `group.prob` (probability of cells belonging to a particular group): (⅓, ⅓, ⅓) everywhere except (.75, .25, .05) for the uneven groups dataset.
- `de.prob` (probability of a genes being differentially expressed across groups): 0.7 for all datasets, except 0.4 for the overlapping clusters dataset.

Counts are log-normalized in the same way as real scRNA-seq data.
